# Impact of second-generation antipsychotics on white matter microstructure in adolescent-onset psychosis

**DOI:** 10.1101/721225

**Authors:** Claudia Barth, Vera Lonning, Tiril Pedersen Gurholt, Ole A. Andreassen, Anne M. Myhre, Ingrid Agartz

## Abstract

White matter abnormalities are well-established in adult patients with psychosis. Yet less is known about changes in early onset psychosis (EOP) during adolescence, especially whether antipsychotic medication might impact white matter microstructure in this sensitive phase. Here, we utilized Magnetic Resonance Imaging (MRI) in unmedicated and medicated adolescent EOP patients in comparison to healthy controls to examine the impact of antipsychotic medication status on indices of white matter microstructure. Twenty-two EOP patients (11 unmedicated) and 33 healthy controls, aged between 12-18 years, underwent 3T diffusion-weighted MRI. Using Tract-based Spatial Statistics, we calculate case-control differences in scalar diffusion measures, i.e. fractional anisotropy (FA), axial diffusion (AD) and radial diffusion (RD), and investigated their association with antipsychotic medication. We found significantly lower mean FA and AD in largely overlapping areas, particularly in left anterior corona radiata (ACR), in EOP patients relative to healthy controls. Mean FA in the left ACR was significantly associated with antipsychotic medication status (t = 2.991, p = 0.008, R^2^ = 0.298), showing higher FA values in medicated compared to unmedicated EOP patients. The present study is the first to link antipsychotic medication status to altered regional FA in the left ACR, a region being discussed to contribute to the etiology of psychosis. Yet, further work with larger samples is needed to draw firm conclusions about putatively enhancing effects of antipsychotic medication on white matter microstructure early in the disease process.

## Introduction

A variety of hypotheses has been proposed to explain the etiology of psychotic disorders, including aberrant dopamine neurotransmission ^1^, altered neurodevelopmental trajectories ^2^, and active neuroinflammation ^3^. Such theories are not mutually exclusive and are more likely complementary. Patients with early onset psychosis (EOP), with defined age of onset before 18 years, provide an unprecedented opportunity to specifically investigate the perspective of aberrant neurodevelopment.

The application of diffusion weighted imaging (DWI) can relate white matter organization to disease. DWI maps the Brownian movement of water molecules in the brain in vivo, and as axon membranes and myelin provide natural barriers for water diffusion, DWI can be used to infer local tissue properties ^4^. The most commonly used scalar measure is fractional anisotropy (FA), which characterizes the degree of diffusion directionality. For an in-depth evaluation of FA, the relative contribution of axial diffusion (AD) along the primary axis, and radial diffusion (RD) perpendicular to it, can be informative. RD has been associated with changes in myelin ^5^, and a disruption of myelin sheaths may be reflected in an increased RD. Conversely, AD has been linked to axonal integrity and axonal damage may be characterized by decreased AD ^4^.

Using DWI, a number of studies showed widespread FA reductions in many different brain regions with low spatial overlap such as corpus callosum, cingulum, superior longitudinal fasciculus, inferior longitudinal fasciculus and fronto-occipital fasciculus in EOP patients compared to healthy controls ^6-10^. Scalar DWI measures beyond FA are rarely analyzed in EOP populations. However, Lagopoulos and colleagues report on increased RD values, indicative of potential demyelinating processes underlying the observed white matter abnormalities ^10^.

The low degree of regional specificity of whiter matter changes seems to be attributed to a number of factors including differences in image acquisition, different analysis approaches (ROI vs voxel-wise), small sample sizes, low prevalence of EOP (estimated prevalence of 17.6 in 10,000 at age of 18 years ^11^) and differing sample characteristics such as age of onset. Further, antipsychotic medication status might also affect the pattern of white matter microstructure in EOP.

Studies investigating white matter microstructure in EOP mainly focus on case-control differences, either reporting antipsychotic effects as secondary findings or using antipsychotic medication status as a covariate of no interest. So far, studies in EOP patients do not indicate an impact of either current ^12-14^ or cumulative antipsychotic exposure ^6,13,15^ on scalar DWI measures. The absence of an antipsychotic medication effect could reflect small sample sizes and young patients with shorter medication histories. The apparent limitations of the adolescent study population also hold potential advantages: EOP patients are less affected by chronic exposure to antipsychotic medication in comparison to their adult counterparts, which allows for dissecting medication-mediated from disease-related effects on brain structure. Furthermore, according to the World Health Organization guideline for pharmacological interventions in adolescents with psychotic disorders (2015), antipsychotic medication use is less recommended in comparison to adult patients with psychosis ^16^. This partly translates to a clinical practice of a higher reluctance in starting antipsychotic treatment early in the course of psychosis in children and adolescents, leading to a higher percentage of antipsychotic-naïve EOP patients relative to adult first episode patients with psychosis. Thus, EOP patients represent an ideal population to investigate the impact of antipsychotic medication on white matter structure early in disease progression.

Here, we use a thoroughly clinically characterized adolescent EOP sample to (1) investigate white matter microstructure in comparison to healthy controls, and (2) explore the association between second-generation antipsychotic medication and white matter microstructure in medicated compared to currently unmedicated/antipsychotic-naïve EOP patients. We utilize DWI and, by using Tract-Based Spatial Statistics (TBSS), we calculate FA and its scalar sub-measures, RD and AD, and investigate their association with antipsychotic medication and other clinical measures (e.g. Positive and Negative Syndrome Scale, etc.). Based on the existing literature, we hypothesized that EOP patients show widespread reduced FA attended by increased RD and unchanged AD compared to healthy controls, mainly in the corpus callosum and superior/inferior longitudinal fasciculus. As there are no established effects of antipsychotic medication on white matter structure in EOP patients, our post hoc analysis is exploratory by nature.

## Results

### Demographic and clinical data

Sample demographics and clinical characteristics separated by antipsychotic medication status of EOP patients are reported in Table 1 and Table 2, respectively.

**Table 1.** General sample characteristics.

|  | *EOP patients overall*<br>*N = 22* | *Healthy controls*<br>*N = 33* | *Statistics*<br>*group-level* |
| --- | --- | --- | --- |
| *Sex, female N (%)* | 15 (68.2) | 20 (60.6) | X<sup>2</sup>, p = 0.775 |
| *Age at MRI (y)* | 16.71 [16.24, 17.57] | 16.19 [15.06, 17.26] | KW, p = 0.128 |
| Range | 14.53 – 18.25 | 12.67 – 18.15 |  |
| *Handedness N (%)* |  |  | FET, p = 0.634 |
| Right | 18 (81.8) | 30 (90.9) |  |
| Left | 2 (9.1) | 2 (6.1) |  |
| Missing N (%) | 2 (9.1) | 1 (3.0) |  |
| *Parental Education (y)* |  |  |  |
| Mother | 15.00 [14.25, 16.00] | 16.00 [15.00, 17.00] | KW, p = 0.106 |
| Range | 11 – 19 | 12 – 22 |  |
| Missing N (%) | 0 | 2 (6.1) |  |
| Father | 15.00 [12.00, 16.00] | 15.50 [15.00, 17.00] | KW, p = 0.133 |
| Range | 10 – 23 | 11 – 20 |  |
| Missing N (%) | 1 (4.5) | 3 (9.1) |  |
| *IQ* | 103.00 [95.50, 109.50] | 102.00 [94.75, 110.75] | KW, p = 0.922 |
| Range | 83 – 132 | 70 – 116 |  |
| Missing N (%) | 3 (13.63) | 1 (3) |  |
| *CGAS* | 45.00 [36.25, 53.75] | 91.00 [83.00, 95.00] | KW, *p < 0.001* |
| Range | 32 – 59 | 75 – 98 |  |
| *MFQ* | 28.00 [18.00, 38.00] | 5.00 [1.00, 8.25] | KW, *p < 0.001* |
| Range | 5 – 52 | 0 – 31 |  |
| Missing N (%) | 1 (4.5) | 1 (3) |  |
| *BMI (kg/m<sup>2</sup>)* | 20.50 [18.00, 23.15] | 20.70 [18.58, 22.18] | KW, p = 0.704 |
| Range | 15.4 – 35.4 | 16.7 – 26.0 |  |
| Missing N (%) | 3 (13.6) | 5 (15.2) |  |
| *Cannabis use, yes N (%)* | 7 (31.8) | 1 (3.0) | FET, **p = 0.005** |
| *Alcohol use, yes N (%)* | 12 (54.5) | 15 (45.5) | X<sup>2</sup>, p = 0.700 |

\*non-normal distributed data in median [interquartile range], N = Number of participants, m = male, f = female, y = years, r = right, l = left, IQ = Intelligence Quotient, BMI = Body Mass Index, CGAS = Children's Global Assessment Scale, MFQ = Mood and Feelings Questionnaire, KW = Kruskal-Wallis Test, FET = Fisher's Exact Test

**Table 2.** Patient clinical characteristics stratified by antipsychotic medication status.

|  | *EOP patients*<br>*Off AP at scan*<br>*N = 11* | *EOP patients*<br>*On AP at scan*<br>*N = 11* | *Statistics*<br>*patient-level* |
| --- | --- | --- | --- |
| *Sex, female N (%)* | 6 (54.5) | 9 (81.8) | FET, $p = 0.361$ |
| *Age Scan (y)* | $15.94 \pm 1.00$ | $17.44 \pm 0.65$ | *t-test, $p < 0.001$* |
| Range | 14.53 – 17.25 | 16.53 – 18.25 |  |
| *BMI (kg/m<sup>2</sup>)* | 19.20 [17.40, 22.90] | 20.55 [18.93, 25.43] | KW, $p = 0.457$ |
| Range | 15.4 – 28.7 | 16.3 – 35.4 |  |
| Missing N (%) | 0 | 3 (27.3) |  |
| *CGAS* | $46.00 \pm 8.00$ | $43.91 \pm 9.38$ | t-test, $p = 0.580$ |
| Range | 34 – 59 | 32 – 58 |  |
| *MFQ* | $30.82 \pm 12.60$ | $27.10 \pm 13.39$ | t-test, $p = 0.520$ |
| Range | 5 – 52 | 8 – 49 |  |
| Missing N (%) | 0 | 1 (9.1) |  |
| *PANSS* |  |  |  |
| *positive* | $19.09 \pm 3.83$ | $17.18 \pm 3.84$ | t-test, $p = 0.257$ |
| Range | 12 – 25 | 13 – 26 |  |
| *negative* | $22.09 \pm 7.13$ | $18.27 \pm 6.96$ | t-test, $p = 0.218$ |
| Range | 9 – 32 | 7 – 32 |  |
| *general* | $38.73 \pm 8.44$ | $36.27 \pm 7.89$ | t-test, $p = 0.489$ |
| Range | 28 – 54 | 24 – 52 |  |
| *Age of Onset (y)* | $14.39 \pm 1.92$ | $14.83 \pm 2.07$ | t-test, $p = 0.614$ |
| Range | 10 – 16 | 12 – 17.6 |  |
| *DUP (w)* | 36.00 [23.00, 82.00] | 12.00 [8.00, 18.00] | KW, $p = 0.005$ |
| Range | 14 – 227 | 3 – 125 |  |
| *DUI (y)* | 0.75 [0.66, 1.79] | 1.76 [0.64, 4.52] | KW, $p = 0.375$ |
| Range | 0.47 – 4.53 | 0.33 – 5.97 |  |
| *Diagnosis* | | | FET, $p = 1.000$ |
| SCZ | 7 | 7 |  |
| SCA | 1 | 0 |  |
| NOS | 3 | 4 |  |
| *Antipsychotics* |  |  |  |
| Aripiprazole |  | 5 |  |
| Risperidone |  | 3 |  |
| Quetiapine |  | 3 |  |
| *CPZ* |  |  |  |
| current | | $272.3 \pm 140.83$ | |
| Range |  | 151.52 – 559.44 |  |
| *cumulative* | 0.08 ± 0.28 | 21.6 ± 19.13 |  |
| (AP-naïve, N = 9) |  |  |  |
| Range | 0 – 0.92 | 1.69 – 59.28 |  |
| **Cannabis use, yes N (%)** | 3 (27.3) | 4 (36.4) | FET, p = 1.000 |
| **Alcohol use, yes N (%)** | 4 (36.4) | 8 (72.7) | FET, p = 0.198 |

\*normally distributed data as mean ± standard deviation, non-normal distributed data in median [interquartile range], AP = antipsychotics, N = number of participants, m = male, f = female, y = years, r = right, l = left, IQ = Intelligence Quotient, BMI = Body Mass Index, CGAS = Children's Global Assessment Scale, MFQ = Mood and Feelings Questionnaire, PANSS = Positive and Negative Symptom Scale, DUP = Duration of Untreated Psychosis, DUI = Duration of illness, SCZ = schizophrenia, SCA = schizoaffective, NOS = psychosis, not other specified, CPZ = chlorpromazine equivalent, KW = Kruskal-Wallis Test, FET = Fisher's Exact Test.

### TBSS analyses

Voxel-wise statistical analysis of case-control differences revealed lower mean FA values in the left genu of the corpus callosum, the left anterior corona radiata (ACR), and the right superior longitudinal fasciculus (SLF) in EOP patients compared to healthy controls (see Figure 1 and Table 3). There was no increase in mean FA for the opposing contrast.

**Table 3.** White matter cluster of reduced fractional anisotropy in early onset psychosis patients relative to healthy controls.

| CIuster | Region*<sub>1</sub> | Side | Voxels | MNI coordinates in mm |  |  | t-values |
| --- | --- | --- | --- | --- | --- | --- | --- |
|  |  |  |  | X | Y | Z |  |
| 4 | Genu of corpus callosum | L | 150 | -14 | 35 | 6 | 3.35 |
| 3 | Anterior corona radiata*<sub>2</sub> | L | 46 | -26 | 13 | 14 | 3.69 |
| 2 | Superior longitudinal fasciculus | R | 12 | 29 | -37 | 35 | 4.81 |
| 1 | Anterior corona radiata (16% Inferior fronto-occipital fasciculus) *<sub>3</sub> | L | 9 | -26 | 24 | 16 | 4.05 |

\*<sub>1</sub> Johns Hopkins University International Consortium for Brain Mapping (JHU ICBM)-DTI-81 white matter atlas and JHU white matter tractography atlas (in brackets) were utilized to label significant clusters with specific tract names
\*<sub>2</sub> 6% uncinate fasciculus/ 5% inferior fronto-occipital fasciculus according to JHU White-Matter Tractography Atlas
\*<sub>3</sub> 11% anterior thalamic radiation, 8% uncinate fasciculus according to JHU White-Matter Tractography Atlas

**Figure 1.**
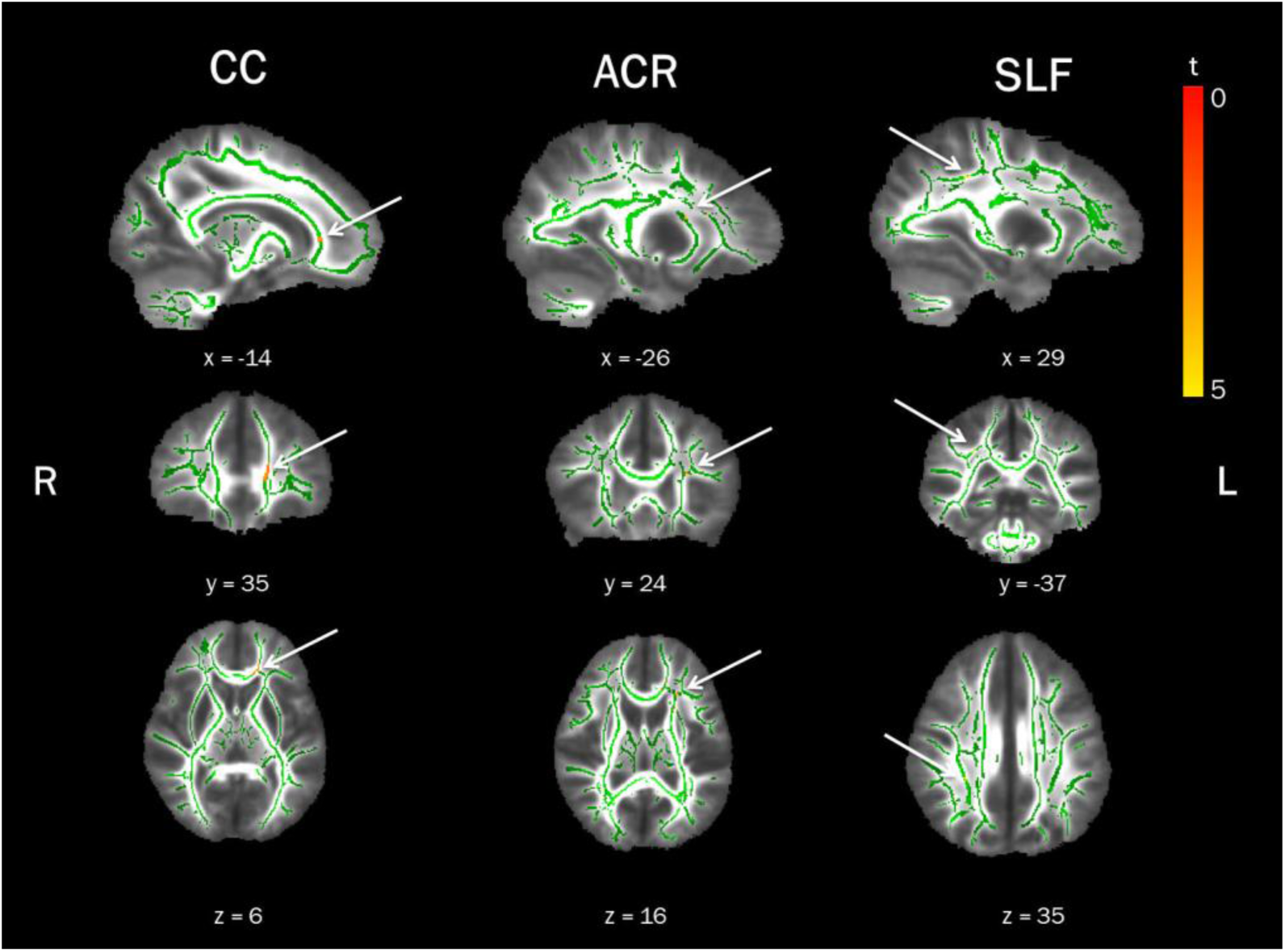
Lower fractional anisotropy (FA) in early onset psychosis (EOP) patients in comparison to healthy controls. Displayed are significant FWE-corrected TBSS results (red-yellow, p ≤ 0.01), contrasting EOP patients against healthy controls, overlaid on the study-specific mean FA skeleton in green and the mean FA image. Results shown underwent threshold-free cluster enhancement and are corrected for age and sex. CC = corpus callosum, ACR = anterior corona radiata, SLF = superior longitudinal fasciculus, R = right, L = left.

Applying the TBSS pipeline to diffusion-derived data other than FA, namely RD and AD, did not yield significant case-control differences for RD, but lower mean AD values overlapping with FA findings in EOP patients in comparison to healthy controls. In detail, mean AD values were significantly lower in the left ACR (Table S2, available online), in addition to lower values in the right posterior limb of the internal capsule (PLIC) and right superior fronto-occipital fasciculus (SFOF).

Extracted mean values of all scalar diffusion measures for all significant clusters stratified by group are displayed in supplementary Figure S1 (available online) for descriptive purposes only. In the interest of transparency, TBSS case-control differences in mean FA and mean AD (patients < controls) at a cluster-forming threshold of p ≤ 0.05 are also presented (see supplementary Figure S2, available online).

### Linear regression analyses

To evaluate the potential influence of duration of illness and antipsychotic treatment on regional mean FA and mean AD values within significant TBSS clusters, in patients, linear regression analyses were performed. While duration of illness was not significantly associated with mean FA in any of the clusters, we found a significant negative association with mean AD in the left ACR (t = -2.364, p = 0.029). For latter association, however, the regression equation was not significant (p = 0.085) and the overall explanatory power of the model was low (R^2^ = 0.147), rendering this finding most likely spurious (see supplementary Table S1, available online).

Exposure to antipsychotic medication was significantly associated with mean FA values in the left ACR (t = 2.991, p = 0.008, R^2^ = 0.298), showing higher mean FA in medicated relative to the unmedicated patients (Figure 2). There was no association between antipsychotic medication and mean FA or mean AD in the other significant TBSS clusters (see supplementary Table S1, available online). Cohen’s d effect size values suggest a high (d = 1.48) and a low (d = -0.08) standardized difference in mean FA of the left ACR in unmedicated and medicated EOP patients relative to healthy controls, respectively.

**Figure 2.**
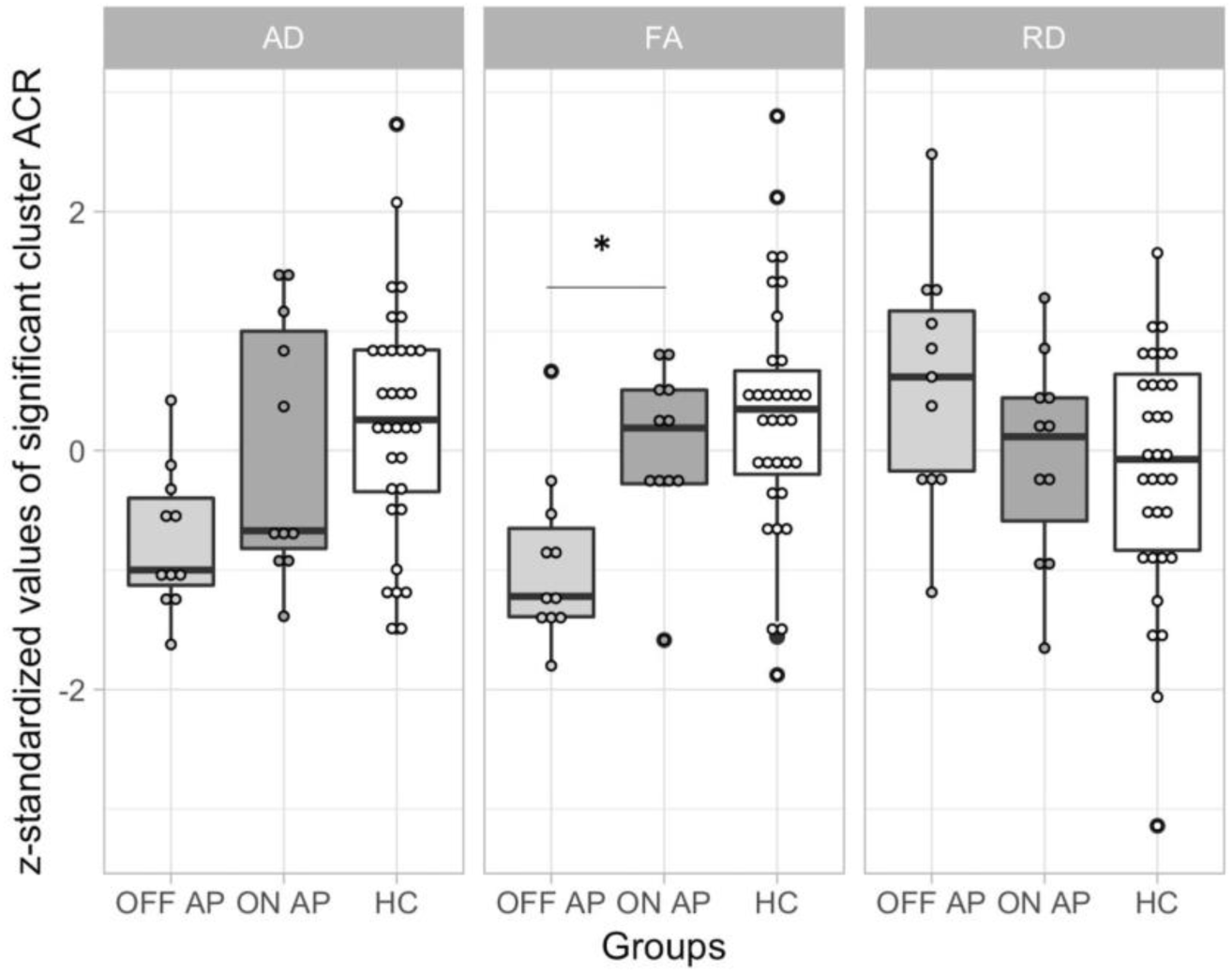
Extracted scalar diffusion values of the left anterior corona radiata (ACR) cluster stratified by antipsychotic use, in comparison to the healthy controls (HC). Data is z-standardized and presented as boxplots for the different scalar diffusion measures overlaid with raw data points. HC are depicted in white, EOP patients on antipsychotic medication in dark grey and EOP patients off antipsychotic medication in light grey. EOP = Early onset psychosis, AP = Antipsychotic use (on = yes, off = no), AD = axial diffusion, FA = fractional anisotropy, RD = radial diffusion. Significant differences in scalar measures between patient subgroups, based on linear regression models, are indicated with a star.

Cohen’s d effect size values suggest a high (d = 1.48) and a low (d = -0.08) standardized difference in mean FA of the left ACR in unmedicated and medicated EOP patients relative to healthy controls, respectively. Based on visual inspection, higher mean FA in medicated patients seems driven by an increase in AD and a decrease in RD (Figure 2). However, extracted mean AD and RD did not differ significantly between medicated and unmedicated patients (Welch Two Sample t-test, AD: t = 0.183, p = 0.857; RD: t = 1.887, p = 0.079).

### Association with clinical measures

We found no association between current antipsychotic medication evaluated as chlorpromazine equivalent at scan day (CPZ, Spearman ρ = 0.13, n = 11, p = 0.695) or cumulative CPZ (Pearson ρ = - 0.15, n = 11, p = 0.665) with regional mean FA values in the left ACR.

In addition, we also found no significant correlations, which survived correction for multiple comparisons (FA cluster: Bonferroni, α = 0.003; AD cluster: Bonferroni, α = 0.004), between neither extracted mean FA nor mean AD values of all significant TBSS clusters and clinical measures such as PANSS (neither positive nor negative), CGAS and MFQ scores.

## Discussion

Our case-control results replicate prominent brain regions with known white matter abnormalities implicated in EOP, namely corpus callosum ^6^, right SLF ^14,17,18^ and the left ACR ^9,10^. Lower FA in these regions seems to occur early in the disease process ^10,18,19^. For instance, Lagopoulos and colleagues found a decrease in FA in the left ACR in both patients with established psychiatric disorder and patients exhibiting sub-syndromal symptoms, aged 14-30 years. Based on these findings, the authors proposed that abnormalities in the left ACR are a putative precursor to the development of a psychiatric condition ^10^.

However, the ACR is a highly heterogeneous structure with three long-range association fiber tracts traversing through it ^10^: anterior thalamic radiation (ATR), inferior fronto-occipital fasciculus (IFOF) and uncinate fasciculus (UF). All three association fibers form connections to the frontal lobe and have been implicated in the pathophysiology of psychiatric disorders ^10,20-22^. In the current study, the left ACR peak voxel shows a 16% probability of IFOF involvement based on the JHU White-Matter Tractography Atlas. The IFOF connects the occipital and temporal lobes with the orbitofrontal cortex as part of the ventral visual and language stream. In particular, the left IFOF seems to subserve language semantics ^23^. Already in 1996, Aloia and colleagues proposed that the disruption of semantic networks have potential implications for the origin of “thought disorder” in schizophrenia ^24^. Adding to this hypothesis, patients with 22q11.2 deletion syndrome, who are genetically at high risk for developing schizophrenia, showed lower FA values in left IFOF ^25^. Furthermore, DeRosse and colleagues found that lower FA proximal to the SLF and corticospinal tract bilaterally, and left IFOF and left inferior longitudinal fasciculus (ILF), were associated with higher levels of psychotic-like experiences in otherwise healthy volunteers ^26^. In early-onset schizophrenia (EOS) patients, lower FA in the left IFOF and the left ILF predicted worse neurocognitive performance ^9^. The authors also detected a shared decrease in FA in the left IFOF among patients with clinical high risk for schizophrenia and patients with established EOS, in comparison to healthy controls. Together, these findings suggest that white matter abnormalities in the left ACR, putatively in the left IFOF, may represent a potential candidate for understanding the etiology of psychosis.

This assumption seems further supported by effects of antipsychotic medication on diffusion metrics in the left ACR. We found that FA values in the left ACR were significantly predicted by antipsychotic medication status, with higher FA values in medicated relative to unmedicated EOP patients. No such association was found with the other brain regions showing significantly decreased FA values. Besides the high Cohen’s d effect size estimate, we found no significant association of regional FA with either current or cumulative antipsychotic exposure. This lack of significant associations is, however, in line with previous studies in EOP patients ^6,12-15^ and likely due to the fairly short medication history in EOP patients compared to their adult-onset counterparts or limited sample size. Hence, the presence of antipsychotic medication rather than the actual dose might induce the observed changes in white matter microstructure.

In medicated patients, increased FA values seem to be driven by an increase in AD and a decrease in RD, relative to their unmedicated counterparts (Figure 2). Thus, FA might be enhanced by antipsychotic medication as a result of both facilitated parallel diffusion (AD, potentially mediated by an increase in axon numbers, and restricted perpendicular diffusion (RD), indicative of changes in myelin.

Converging evidence from multiple studies suggests oligodendroglial dysfunction, with subsequent abnormalities in myelin maintenance and repair, to underpin white matter abnormalities observed in psychotic patients ^27^. In the framework of schizophrenia, it has been proposed that myelin dysfunction, especially in frontal regions, contributes to psychotic symptoms ^13,27^. Based on findings from cell culture studies using aripiprazole ^28^ and rodent work using quetiapine ^29,30^, second-generation antipsychotic medication may promote oligodendrocyte recovery and myelin repair leading to reduced white matter abnormalities and, subsequently, reduced psychotic symptoms. A recent study in patients with schizophrenia also reports on promyelinating effects of antipsychotics ^31^. Tishler and colleagues found an increase in intracortical myelin predominantly mediated by risperidone and other second-generation antipsychotics in adult patients with schizophrenia compared to healthy controls within the first year of treatment. In the current study, given that medicated EOP patients either received aripiprazole, quetiapine or risperidone, one might speculate that early medication with second-generation antipsychotics might affect white matter microstructure by remediating oligodendroglial dysfunction, leading to an increase in FA detected by DWI.

Even though FA is highly sensitive to microstructural changes in general, it lacks neurobiological specificity to the exact type of change ^4^. For instance, a decrease in FA can reflect alternations in fiber organization, including packing density and fiber crossing, and myelin loss or myelin remodeling ^32^. The interpretation of FA and its scalar sub-measures is further complicated by the lack of sensitivity in regions of crossing white matter tracts, such as the ACR. Yet, as stated by Alexander and colleagues ^4^, this confound is unavoidable as many areas of the brain have considerable areas of fiber crossing. Here, we found a widespread decrease of AD on a whole brain level, indicative of axonal damage, but no changes were found in RD, relative to healthy controls. This finding is not in line with previous work from Lagopoulos and colleagues, who found a decrease in FA in the left ACR associated with increases in RD and no changes in AD ^10^.

Although there are likely several reasons for these conflicting findings, neuroinflammatory processes might pose particular difficulties in interpreting DWI signals in psychotic populations ^33^. For instance, in an animal model of cuprizone-induced demyelination of corpus callosum, regions with extensive axonal edema and prominent cellular inflammation showed no change in RD, while AD values were diminished at the beginning of demyelination ^34^. Given the neuroinflammation hypothesis of schizophrenia ^3^, it seems likely that the disease progression encompasses a dynamic evolution of inflammation, axonal injury, and myelin degeneration. In the current study, one might speculate that EOP patients are on the verge of undergoing demyelination processes, reflected by widespread decreases in AD. However, the timing of neuroinflammation in psychotic disorders relative to tissue injury is unclear, leading to a heightened risk of misinterpreting changes in DWI measures. According to a recent review of Winklewski and colleagues, in cases of neuroinflammation linked to tissue damage, DWI seems to underestimate the extent of demyelination (undervalued RD), and overestimate the extent of axonal injury (overvalued AD) ^35^. This pattern seems replicated in our study, with significant changes in AD and no changes in RD. As the consistency of DWI metrics seems affected by brain edema and inflammatory response, future studies can benefit from using tools such as free water imaging which provide the opportunity to separate the contribution of extracellular water from the diffusion of water molecules inside the fiber tracts, leading to a higher specificity in detecting structural changes ^36^.

Deviations in scalar DWI measures in the current study relative to previous studies could also be due to ongoing white matter maturation processes in our adolescent EOP sample. In healthy individuals, age-related increases in FA during childhood, adolescence and early adulthood have been consistently reported ^37-40^. This increase in FA seems primarily driven by a reduction in RD, while AD remains fairly stable or decreases slightly ^41,42^. Findings for AD changes during the transition to adulthood are less consistent ^37-40^. Thus, the AD difference found in the current study could also be attributed to developmental processes, which may fade as adolescents mature into adulthood.

However, it should be noted that the neurodevelopmental trajectories of white matter structure relative to disease progression in EOP patients are unclear. So far, three different studies yielded inconclusive results, either postulating diverging ^13^, converging ^43^, or parallel ^44^ trajectories relative to healthy controls. In the current study, we did not find any predictive value of duration of illness for regional FA. This is in line with previous findings from Kumra and colleagues, who speculated that lower FA in EOP patients compared to healthy controls reflects developmental abnormalities rather than secondary effects of the disease progression ^13^. In addition, Epstein and Kumra found lower FA in the inferior longitudinal fasciculus, IFOF and corticospinal tract, but no significant group differences in longitudinal changes in FA ^44^. Thus, the observed changes in the current study might persist but do not affect the overall white matter maturation trajectories. However, the cross-sectional nature of the current study precludes the assessment of developmental effects over time.

The results of the current study should be considered in the context of several limitations. Unmedicated EOP patients were significantly younger than those receiving medication. Although the analysis was corrected for age, we cannot exclude that the age of the patients contributes to the observed antipsychotic-medication related changes in white matter structure. It is possible that time-of-measurement effects, with older patients having higher FA values than younger patients due to more advanced white matter maturation, could confound our results. However, while we acknowledge this possibility, we consider this unlikely as we would expect differences in FA values in other regions showing a similar maturation trajectory, such as the SLF ^41^, if our results were mainly driven by age differences. This was not the case, as we found no significant difference in mean FA values of the SLF between medicated and unmedicated patients (t = -0.56934, p-value = 0.576).

We are also aware of the limitations of our small sample size. While we acknowledge the possibility that our results might be spurious and limited to our EOP sample, we consider this unlikely to be the driving factor for the following reasons: (1) using a stringent p-value of 0.01 to only accept an error rate of 1% for the whole-brain TBSS (and subsequent mean FA extractions), we replicated white matter microstructure abnormalities in brain regions implicated in the pathophysiology of EOP and adult-onset schizophrenia; and (2) in the current study, patients on second-generation antipsychotic medication received either aripiprazole, quetiapine or risperidone, drugs which have previously been associated with oligodendrocyte recovery and myelin repair in cell-culture ^28^, rodent ^29,30^ and human studies ^31^. In summary, a very conservative p-threshold and findings from the literature and our own data support that second-generation antipsychotic medication might impact white matter microstructure in schizophrenia. To the best of our knowledge, our study is the first to highlight this potential association in a well-characterized and balanced sample of medicated and unmedicated EOP patients. Yet, a replication of the results in a bigger sample is warranted.

In summary, the present study is the first to link antipsychotic medication status to altered regional FA in the left ACR in patients with EOP. Understanding the significance of white matter abnormalities in the left ACR in adolescents with EOP and the putatively remediating effect of antipsychotic medication, may help to phenotype the disease and to develop new pharmacological regimes to subsequently improve functional outcome. Assuming that antipsychotic medication reverses the hypothesized myelin dysfunction in psychosis, early interventions with antipsychotic medication, already in individuals at risk of developing psychosis, could provide the opportunity to normalize white matter maturation. Although exciting, further work is needed to draw firm conclusions about the putative enhancing effects of antipsychotic medication early in the disease process. Building on our first results, longitudinal studies with larger samples sizes using high resolution DWI in combination with clinical, genetic and neurocognitive measures are warranted to delineate heritability, affected brain regions, antipsychotic medication effects, and directions of FA changes over time.

## Methods

### Participants

The study sample was drawn from the ongoing longitudinal Youth-Thematic-Organized-Psychosis (Youth-TOP) research study, which is a subdivision of the TOP research group/NORMENT and KG Jebsen center of psychosis research in Oslo, Norway. EOP patients, aged between 12-18 years, were recruited from in- and outpatient clinics in the Oslo region. Healthy controls were randomly selected from the Norwegian National Registry in the same catchment areas. All participants and their respective parents/guardians provided written informed consent. The study was approved by the Regional Committee for Medical Research Ethics (REK-Sør) and the Norwegian Data Inspectorate and was conducted in accordance with the Declaration of Helsinki.

For study inclusion, participants were required to have an intelligence quotient (IQ) > 70, a good command of the Norwegian language, no previous moderate to severe head injuries, no diagnosis of substance-induced psychotic disorder, and no organic brain disease. IQ was measured by the Wechsler Abbreviated Scale of Intelligence. Diagnosis was established according to the Diagnostic and Statistical Manual of Mental Disorder-IV criteria using the Norwegian version of the Kiddie-Schedule for Affective Disorders and Schizophrenia for School Aged Children (6-18 years): Present and Lifetime Version (K-SADS-PL). The clinical characterization was conducted by trained psychologists or psychiatrists.

A total of 67 participants (27 patients/40 controls) satisfied the above-mentioned criteria and underwent MRI examination. All MRI scans were visually inspected by a trained neuroradiologist to rule out any pathological changes. Out of the initial sample, seven control participants and five patients were excluded due to (i) clinical/radiological reasons (five patients/ three controls), or (ii) strong motion artefacts in the diffusion imaging data (four controls), resulting in a final sample of 55 participants (22 patients/ 33 controls) being entered in the statistical analysis.

### Clinical measures

Presence and severity of psychopathological symptoms of EOP patients were assessed using the Positive and Negative Syndrome Scale (PANSS). Children Global Assessment Scale (CGAS) and Mood and Feelings Questionnaire (MFQ, long version) were evaluated in all participants to measure general functioning level and to screen for depressive symptoms, respectively. Recreational drug use was assessed within the structured K-SADS interview and scored with 0 or 1 for absent or present. For EOP patients, current and lifetime cumulative use of medication was recorded and converted into chlorpromazine equivalents (CPZ), using formulas published elsewhere ^45^. While 11 EOP patients were off any antipsychotic medication at scan, yielding a lack of current CPZ values, three patients had received pharmacological treatment prior to inclusion, resulting in a low cumulative CPZ dosage for this subgroup (see Table 2).

### MRI data acquisition

MR images were acquired on a 3-Tesla General Electric Signa HDxt scanner equipped with an 8-channel head coil at the Oslo University Hospital, Norway. The diffusion imaging data was acquired using a 2D spin-echo whole-brain echo-planar imaging sequence with the following parameters: slice thickness = 2.5 mm, repetition time = 15 s, echo time = 85 ms, flip angle = 90°, acquisition matrix = 96 × 96, in-plane resolution = 1.875 × 1.875 mm. A total of 32 volumes with different gradient directions (b = 1000 s/mm^2^), including two b0-volumes with reversed phase-encode (blip up/down), were acquired.

### Diffusion data analysis

Diffusion data were analyzed with FSL version 5.0.9 using the FMRIB’s software library (https://fsl.fmrib.ox.ac.uk/fsl/fslwiki). Before creating voxel wise maps of diffusion parameters, the following steps of the standard processing pipeline were used: (i) *topup* to correct for susceptibility-induced distortions ^46^, (ii) *eddy* current correction to correct for gradient-coil distortions and head motion ^47^, (iii) removal of non-brain tissue using the Brain Extraction Tool (*bet*) ^48^, and (iv) local fitting of the diffusion tensor at each voxel using *dtifit* (FMRIB’s Diffusion Toolbox (FDT) ^49^). *Dtifit* yielded in voxel wise participant-specific maps of FA, mean diffusion (MD), and axial diffusion (AD, derived from eigenvector λ1). Based on the outputted eigenvectors λ2 and λ3, radial diffusion (RD) was computed ((λ2+λ3)/2)). Next, voxel wise statistical analysis of the FA data was carried out using TBSS ^50^. First, all FA images were nonlinearly aligned to the most representative FA image out of all images and transformed into 1×1×1 mm^3^ MNI152 standard space by means of affine registration. Secondly, TBSS projects all participant’s FA data onto a mean FA tract skeleton (threshold FA > 0.25), before applying voxel wise cross-participant statistics. After TBSS for FA was completed, results were used to generate skeletonized RD and AD data for additional voxel-wise group comparisons using the TBSS non-FA pipeline.

### Statistical analyses

For contrasting case-control differences, we run voxel-wise statistics, co-varied for age and gender, using a nonparametric permutation-based approach (*Randomise*, implemented in FSL, 5000 permutations). All variables were demeaned. The statistical threshold was set at p ≤ 0.01, after family-wise error correction for multiple comparisons using threshold-free cluster enhancement. We chose a highly conservative threshold for FA to minimize type I errors and to better account for the exploratory nature of the study concerning the impact of antipsychotic medication status. For RD and AD, the same statistical model was used.

Regions identified with TBSS (FA) and TBSS non-FA (RD/AD) were subsequently used as masks to extract mean FA, RD and AD values for plotting and further analysis. We refrained from using MD values, as a measure of overall diffusivity within a voxel, in further analysis due to its lack of specificity ^4^. As scalar diffusion measures largely vary in their value ranges, extracted mean values were z-standardized for plotting purposes using the following formula: z = (participant’s value – group mean) / standard deviation.

A linear regression model was performed to examine whether patients’ mean values of significant TBSS and TBSS non-FA clusters were associated with duration of illness and antipsychotic medication status as categorical variable (coded as yes (1)/no (0)). The effect size was reported as Cohen’s d.

If there is an association between patients’ regional mean values and antipsychotic medication, follow-up correlation analysis with current and cumulative CPZ were performed using Spearman’s rank correlation rho for non-normal data.

Further analysis of regional mean values and its association with clinical measures (PANSS, CGAS, and MFQ) were performed using Pearson’s product moment correlation coefficient.

Statistical tests were conducted in R, version 3.5.2 (www.r-project.org).

## Acknowledgements

We thank the study participants and the Youth-TOP clinicians involved in recruitment and assessment at the Norwegian Centre for Mental Disorders (NORMENT) and the Diakonhjemmet Hospital, Oslo, Norway (Runar Elle Smelror, Kirsten Wedervang-Resell, Cecilie Haggag Johannessen, Tarje Tinderholt, Tove Matzen Drachmann). Further, we like to thank Kristine Engen and Brian Frank O’Donnell for proofreading. This work was supported by the Research Council of Norway, grant numbers 223273, 213700, and 250358; the South-Eastern Norway Regional Health Authority, grant numbers 2016-118 and 2017-097; and KG Jebsen Centre for Psychosis Research.

## Author Contributions

CB undertook the processing of the imaging data, the statistical analysis, the literature search, interpreted the results and wrote the first draft of the manuscript. TPG helped with the processing of the imaging data. VL was significantly involved in the participant inclusion and data acquisition for Youth-TOP and calculated current and cumulative chlorpromazine equivalents. IA designed the ongoing longitudinal Youth-TOP research study the data is drawn from. IA, AMM and OAA obtained funding and contributed to the data acquisition. All authors contributed to the critical revision of the manuscript and approved the final draft for submission.

