## Supplementary Materials for "Impact of second-generation antipsychotics on white matter microstructure in adolescent-onset psychosis"

**Supplementary information**

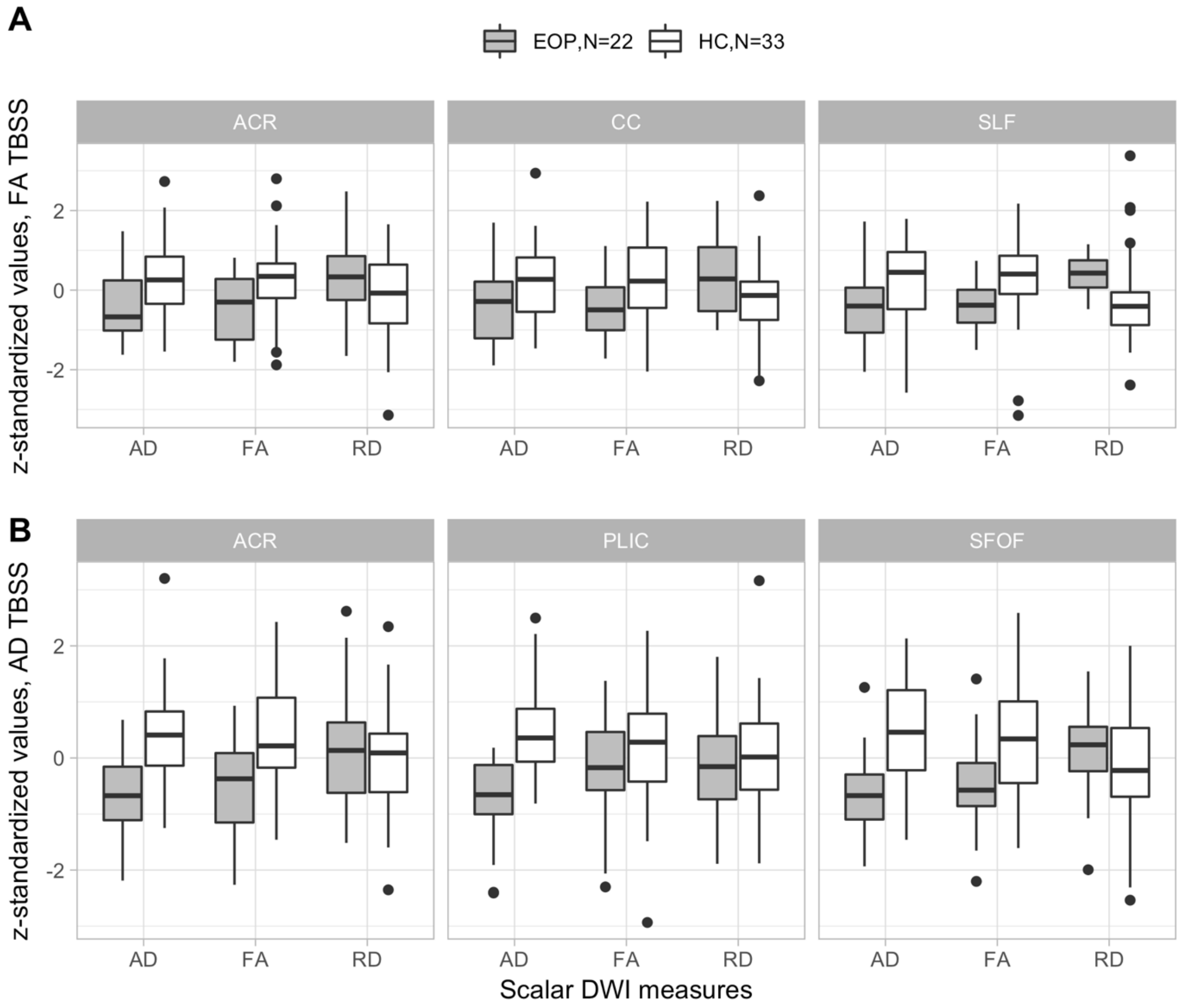


**Figure S1| Extracted mean fractional anisotropy (FA), axial diffusivity (AD) and radial diffusivity (RD) values of significant FA (A) and AD clusters (B) identified with whole-brain TBSS.** Data is presented as grey boxplots for early onset psychosis (EOP) patients and white boxplots for healthy controls (HC). ACR = anterior corona radiata, CC = corpus callosum, SLF = superior longitudinal fasciculus, PLIC = Posterior limb of the internal capsule, SFOF = superior fronto-occipital fasciculus. Note: Data is presented for descriptive purpose only.

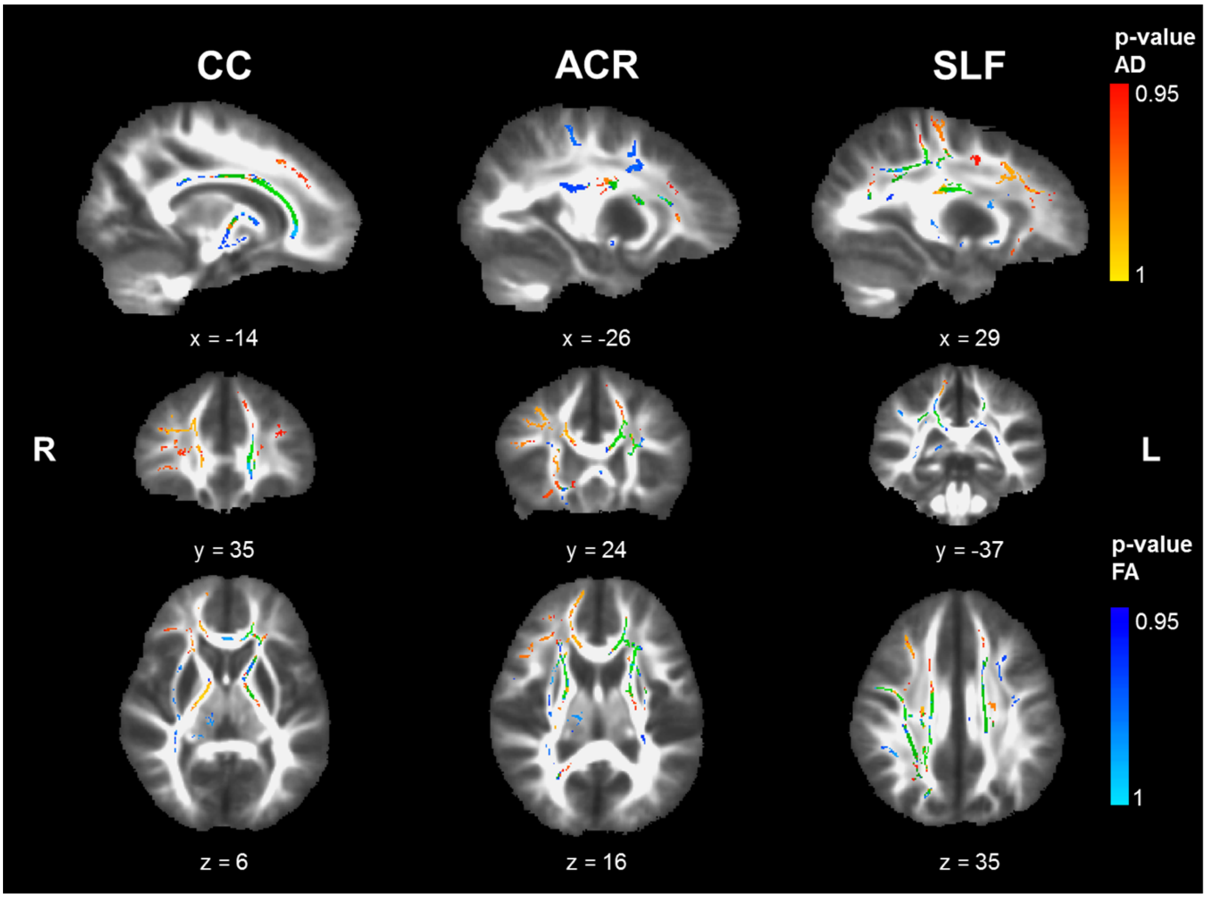


**Figure S2| Overlap of decreased fractional anisotropy (FA) and axial diffusivity (AD) in early onset psychosis (EOP) patients in comparison to healthy controls.** Displayed are significant FWE-corrected TBSS results for FA (light blue – dark blue, $p \leq 0.05$) and AD (red-yellow, $p \leq 0.05$), contrasting EOP patients against healthy controls, overlaid on the study-specific mean FA image. The overlap of both DWI measures is depicted in green. Results shown underwent threshold-free cluster enhancement and are corrected for age and sex. CC = corpus callosum, ACR = anterior corona radiata, SLF = superior longitudinal fasciculus, R = right, L = left. Note: Data is presented for descriptive purpose only.

### Tables

**Table S1| Results of linear regression models with extracted mean fractional anisotropy (FA) and mean axial diffusivity (AD) values of significant TBSS clusters.**

| Model *[1] | Estimate | S.E. | t value | p value | F | df | p | *Adj.* $R^2$ |
|---|---|---|---|---|---|---|---|---|
| **FA TBSS** | | | | | | | | |
| **ACR-IFOF** | | | | | 5.5 | 19 | 0.013 | 0.298 |
| DUI | 0.002 | 0.004 | 0.479 | 0.637 | | | | |
| AP | 0.048 | 0.016 | 2.991 | **0.008** | | | | |
| **SLF** | | | | | | | | |
| DUI | 0.009 | 0.008 | 1.190 | 0.249 | 0.9 | 19 | 0.434 | -0.012 |
| AP | 0.006 | 0.028 | 0.205 | 0.840 | | | | |
| **ACR** | | | | | | | | |
| DUI | 0.0001 | 0.006 | 0.020 | 0.985 | 1.0 | 19 | 0.388 | -0.001 |
| AP | 0.027 | 0.020 | 1.344 | 0.195 | | | | |
| **CC** | | | | | | | | |
| DUI | 0.001 | 0.005 | 0.210 | 0.836 | 0.02 | 19 | 0.977 | -0.103 |
| AP | -0.0003 | 0.016 | -0.016 | 0.987 | | | | |
| **AD TBSS** | | | | | | | | |
| **ACR-ATR** | | | | | | | | |
| DUI | -1.2e-05 | 5.1e-06 | -2.364 | **0.029** | 2.81 | 19 | 0.085 | 0.147 |
| AP | 1.6e-05 | 1.8e-05 | 0.855 | 0.403 | | | | |
| **PLIC** | | | | | | | | |
| DUI | -1.1e-06 | 5.1e-06 | -0.209 | 0.837 | 0.1 | 19 | 0.942 | -0.098 |
| AP | 6.1e-06 | 1.9e-05 | 0.324 | 0.750 | | | | |
| **SFOF** | | | | | | | | |
| DUI | -4.3e-06 | 5.3e-06 | -0.809 | 0.429 | 0.4 | 19 | 0.6801 | -0.061 |
| AP | -2.2e-06 | 1.9e-05 | -0.113 | 0.911 | | | | |

*[1] Regional mean FA of significant clusters identified with Tract-Based Spatial Statistics; DUI = Duration of illness & AP = Antipsychotic use (coded yes (1)/no (0)); ACR-IFOF = anterior corona radiata, 16% inferior fronto-occipital fasciculus, SLF = superior longitudinal fasciculus, ACR = anterior corona radiata, CC = corpus callosum.

**Table S2| White matter cluster of reduced axial anisotropy in early onset psychosis patients relative to healthy controls.**

| | | | | MNI coordinates in mm | | | |
|---|---|---|---|---|---|---|---|
| **Cluster** | **Region*** | **Side** | **Voxels** | **X** | **Y** | **Z** | **t-values** |
| 3 | Anterior corona radiata (50% Anterior thalamic radiation) | L | 620 | -23 | 18 | 15 | 4.93 |
| 2 | Posterior limb of internal capsule | R | 344 | 16 | -10 | 2 | 5.05 |
| 1 | Superior fronto-occipital fasciculus (18% Anterior thalamic radiation) | R | 138 | 21 | 12 | 22 | 5.17 |

* Johns Hopkins University International Consortium for Brain Mapping (JHU ICBM)-DTI-81 white matter atlas and JHU white matter tractography atlas (in brackets) were utilized to label significant clusters with specific tract names
